# *PISD* is a mitochondrial disease gene causing skeletal dysplasia, cataracts, and white matter changes

**DOI:** 10.1101/413070

**Authors:** Tian Zhao, Caitlin M. Goedhart, Pingdewinde Sam, Susanne Lingrell, Adam J. Cornish, Ryan E Lamont, Francois P Bernier, David Sinasac, Jillian S Parboosingh, Care4Rare Canada Consortium, Jean E. Vance, Steven M. Claypool, A Micheil Innes, Timothy E Shutt

**Affiliations:** Alberta Children’s Hospital Research Institute, Department of Medical Genetics, Cumming School of Medicine, University of Calgary, Calgary, Alberta, Canada; Department of Biochemistry & Molecular Biology, Cumming School of Medicine, University of Calgary, Calgary, Alberta, Canada; Department of Physiology, The Johns Hopkins University School of Medicine, Baltimore, MD, USA; Dept of Medicine and Group on Molecular and Cell Biology of Lipids, University of Alberta, Edmonton, Alberta, Canada

## Abstract

Exome sequencing of two sisters with congenital cataracts, short stature and white matter changes identified compound heterozygous variants in the *PISD* gene, encoding the phosphatidylserine decarboxylase enzyme that converts phosphatidylserine (PS) to phosphatidylethanolamine (PE) in the inner mitochondrial membrane (IMM). Decreased conversion of PS to PE, and depletion of total cellular PE levels in patient fibroblasts are consistent with impaired PISD enzyme activity. Meanwhile, as evidence for mitochondrial dysfunction, patient fibroblasts exhibited more fragmented mitochondrial networks, enlarged lysosomes, decreased maximal oxygen consumption rates and increased sensitivity to 2-deoxyglucose. Moreover, treatment with lyso-PE, which can replenish the mitochondrial pool of PE, restored mitochondrial and lysosome morphology in patient fibroblasts. Functional characterization of the *PISD* mutations demonstrates that the maternal variant causes an alternative splice product. Meanwhile, the paternal variant impairs autocatalytic self-processing of the PISD protein required for its activity. Finally, evidence for impaired activity of mitochondrial IMM proteases explains why the phenotypes of these *PISD* patients resemble recently described “mitochondrial chaperonopathies”. Collectively, these findings demonstrate that *PISD* is a novel mitochondrial disease gene.

## Introduction

Mitochondria are double-membrane bound organelles, which in addition to generating most of a cell’s energy via oxidative phosphorylation, also have important roles in regulating many other cellular processes (e.g. apoptosis, immune response, and numerous metabolic pathways). While mitochondrial dysfunction has been implicated in a growing list of human diseases, more severe forms of mitochondrial dysfunction cause a group of rare disorders known as mitochondrial diseases, estimated at ~1 in 4,300 in adults (Gorman, Schaefer et al., 2015). Classic mitochondrial disease is caused by impaired energy production, and often manifests in tissues with high energy demands, such as heart, muscle, brain, and eyes. However, diagnosing mitochondrial disease is difficult due to the clinical and genetic heterogeneity of this group of disorders.

More recently, an atypical class of mitochondrial diseases has been described where impaired mitochondrial protein homeostasis appears to be the underlying cause of mitochondrial dysfunction (Royer-Bertrand, Castillo-Taucher et al., 2015). These ‘ mitochondrial chaperonopathies’ are characterized by atypical skeletal phenotypes and craniofacial features that are not commonly seen in classic mitochondrial disease, as well as cataracts and central nervous system involvement, which are found in mitochondrial disease. To date, only three genes (*LONP1*, *HSPA9*, and *AIFM1*) encoding mitochondrial proteins have been linked to skeletal abnormalities (Dikoglu, Alfaiz et al., 2015, Mierzewska, Rydzanicz et al., 2016, Royer-Bertrand et al., 2015, Strauss, Jinks et al., 2015). The molecular mechanisms through which impaired mitochondrial protein homeostasis lead to skeletal abnormalities, rather than more traditional mitochondrial disease phenotypes, remains unknown.

Maintenance of mitochondrial protein homeostasis is thus a key aspect regulating mitochondrial function, and its impairment leads to disease. Notably, many mitochondrial-specific proteases are bound to the IMM (Quiros, Langer et al., 2015). Thus, it is not surprising that the IMM lipid composition is an important regulator of mitochondrial function (Lu & Claypool, 2015). The *PISD* gene encodes a mitochondrial-localized enzyme that converts phosphatidylserine (PS) to phosphatidylethanolamine (PE) in the inner mitochondrial membrane (IMM). PE, which comprises ~15-25% of cellular membranes, is an important lipid that provides membrane curvature (Vance & Tasseva, 2013). While complete loss of *PISD* is embryonic lethal in mice, highlighting the importance mitochondrial PE, heterozygous mice do not have any overt phenotypes (Steenbergen, Nanowski et al., 2005). In cellular models, severe depletion or complete loss of PISD results in decreased mitochondrial oxidative phosphorylation, and fragmentation of the mitochondrial network (Steenbergen et al., 2005, Tasseva, Bai et al., 2013). Notably, an autocatalytic processing event that generates two subunits (alpha and beta), is required to form a functional PISD enzyme (Li & Dowhan, 1988).

In the present study, we report the first example of patients with pathogenic variants in *PISD*, who presented with congenital cataracts, short stature, mid-face hypoplasia, hypomyelination, ataxia, and intellectual disability. These phenotypes are reminiscent of the skeletal abnormalities described for mutations in *LONP1* (CODAS syndrome), *HSPA9* (EVEN-PLUS syndrome) and *AIFM1* (SEMD), rather than classic mitochondrial disease. Our findings show that mitochondrial protein homeostasis is impaired in fibroblasts from patients with PISD variants. As such, we suggest that *PISD* be included in the list genes associated with impaired mitochondrial protein homeostasis.

## Results

### Clinical data

#### Affected Individual 1 (II-1) (Fig 1)

This individual is now 28 years old. She was born at 36 weeks gestational age after an uneventful pregnancy. Congenital cataracts were diagnosed in the first few months of life and were extracted surgically. At that time the child was generally well, with a normal height and a basic metabolic workup (including screening for galactosemia), which was essentially normal. However, fall off in growth percentiles began in the first year of life. When reassessed at 3 years of age there was evidence of mild developmental delay and recurrent respiratory infections and height was now at −4 to −5 standard deviations below the mean. A skeletal survey done at 8 years of age was non-diagnostic with findings including brachydactyly, delayed bone age, mild thoracic platyspondyly and metaphyseal striations not suggestive of a primary skeletal dysplasia but suggestive of bone hypoproliferation in keeping with an underlying genetic or metabolic syndrome. Increasing difficulty in school became evident, and subsequent psychometric testing was consistent with either mild intellectual disability or borderline intelligence. Serial MRI scans revealed diffuse T2 signal intensity throughout the bihemispheric white matter, with progressive volume loss and hypomyelination of the corpus callosum.

**Figure 1.**
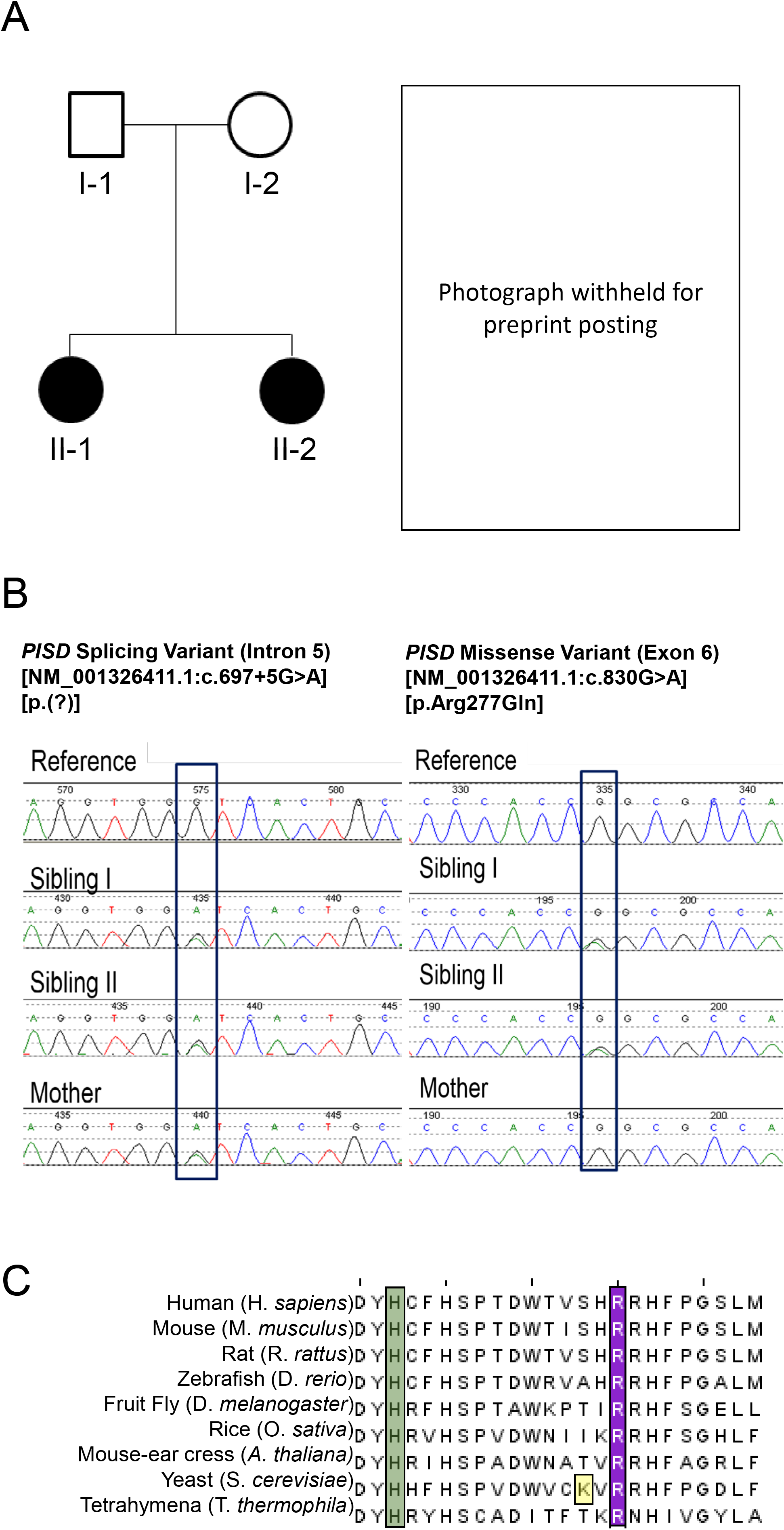
Clinical and genetic patient data. **A)** Pedigree of the patient family along with a picture of the two siblings in infancy. Note strabismus, midface hypoplasia and depressed nasal bridge **B)** Electropherogram conformation of the identified mutations using Sanger sequencing, with mutated residues boxed. **C)** Sequence alignment of PISD homologs from the indicated species showing region containing the R277Q mutation, with the arginine 277 residue highlighted in purple. A conserved histidine residue essential for autocatalysis is highlighted in green and one of four missense mutations in a yeast Psd1p *temperature sensitive* allele is highlighted in yellow (Birner et al., 2003, Choi, Duraisingh et al., 2015, Ogunbona et al., 2017).

She was diagnosed with tracheal stenosis and required a temporary tracheostomy from age 10 to 11 in the context of an acute deterioration with an infectious illness. She was diagnosed at age 18 with bilateral progressive hearing loss. Her final adult height is 124.7cm (−5.3 SD). She had a distinctive facial appearance with depressed nasal ridge and midface hypoplasia (Fig 1A).

#### Affected Individual 2 (II-2)

This is the younger sister of II-1, and these are the only two children born to healthy non-consanguineous parents. She was also diagnosed with congenital cataracts in early infancy. Her early trajectory followed a remarkably similar course to that of her older sister. She also had multiple respiratory illnesses, some requiring ICU admission, mild global developmental delay and progressive short stature. In addition, she developed an intention tremor in early childhood. She has a history of anxiety. She has subglottic stenosis and tracheomalacia and had an ICU admission for a respiratory decompensation at age 25. She was also recently diagnosed at age 25 with mild bilateral sensorineural hearing loss. Skeletal survey and MRI findings were similar to those in her older sister. Her final adult height is 123.7 cm (−5.4 SD). She had midface hypoplasia and a depressed nasal ridge, similar to her older sister (Fig 1A).

The presence of a remarkably similar and striking phenotype in two sisters born to unaffected parents suggested a likely autosomal recessive disorder. The findings including white matter changes on MRI scan, congenital cataracts and progressive hearing loss were potentially suggestive of a progressive neurogenetic or neurometabolic condition. However, screening metabolic investigations had been normal. Clinical diagnoses on the differential diagnosis that all appeared unlikely included Sjogren-Larsson Syndrome, Conradi-Hunnermann syndrome, Hypomyelination-Cataract syndrome and Cockayne syndrome. The possibility, albeit unlikely, that these two siblings both shared more than one rare genetic condition was also considered; however, it was deemed most likely that they had a novel, previously undescribed genetic condition.

### Variant Analysis

Whole exome sequencing (WES) was performed on DNA extracted from peripheral blood provided by both sisters and their mother. Assuming an autosomal recessive inheritance pattern, we searched for rare coding homozygous or compound heterozygous variants (ExAC minor allele frequency less than 1% and observed in five or fewer other Care4Rare exome projects) that were shared by the sisters. The sisters did not share any rare homozygous variants, but they did share two different rare heterozygous variants in the genes *PISD*, *Actin related protein T1 (ACTRT1)* and *SON DNA-Binding Protein (SON)*. However, the mother carried a single variant, NM_001326411.1(PISD):c.697+5G>A [p.(?)], in only one of the three genes. This confirmed that the sisters’ shared variants in *PISD*, NM_001326411.1(PISD):c.830G>A [p.Arg277Gln] and NM_001326411.1(PISD):c.697+5G>A [p.(?)], were bi-allelic (Fig 1B) and that the remaining variants in ACTRT1 and SON were most likely inherited in cis.

The missense variant, c.830G>A [p.Arg277Gln], altered a highly conserved nucleotide (phyloP: 5.69 [-14.1; 6.4]) and amino acid (GERP score: 5.19) in exon six of *PISD* (Fig 1C). The variant had a CADD score of 35, and was predicted to be tolerated by SIFT (score: 0.19, median: 3.05), disease-causing by MutationTaster (p-value: 1) and probably damaging by PolyPhen [score of 0.920 (sensitivity: 0.68; specificity: 0.90)]. The missense variant was reported with a low frequency of: 0.01923% in Genome Aggregation Database (gnomAD); 0.03% for European American populations in the Exome Sequencing Project (ESP) database; and, 0.02% in 1000 genomes database (accessed May 14^th^, 2018). Meanwhile, the splice variant, c.697+5G>A, was not reported in ClinVar and no individuals were reported to be homozygous for the variant in gnomAD (accessed May 14^th^, 2018). The splice variant was predicted to weaken the splice donor site located at the end of exon five and was not reported in gnomAD, ESP, 1000 genomes, or ClinVar databases (accessed May 14^th^, 2018).

### Fibroblast Characterization

Given that there were no previous reports of individuals with pathogenic variants in *PISD*, combined with the unusual clinical presentation for a mitochondrial disease, we sought to perform additional functional analysis of patient fibroblast cells in order to determine whether mitochondrial and/or PISD function were impaired. To begin, we measured the ability of both patient and age-matched control fibroblast cells to convert PS to PE. We observed an approximately 50% decrease in PE synthesis from PS in the patient fibroblast cells, consistent with impaired PISD activity (Fig 2A).

**Figure 2:**
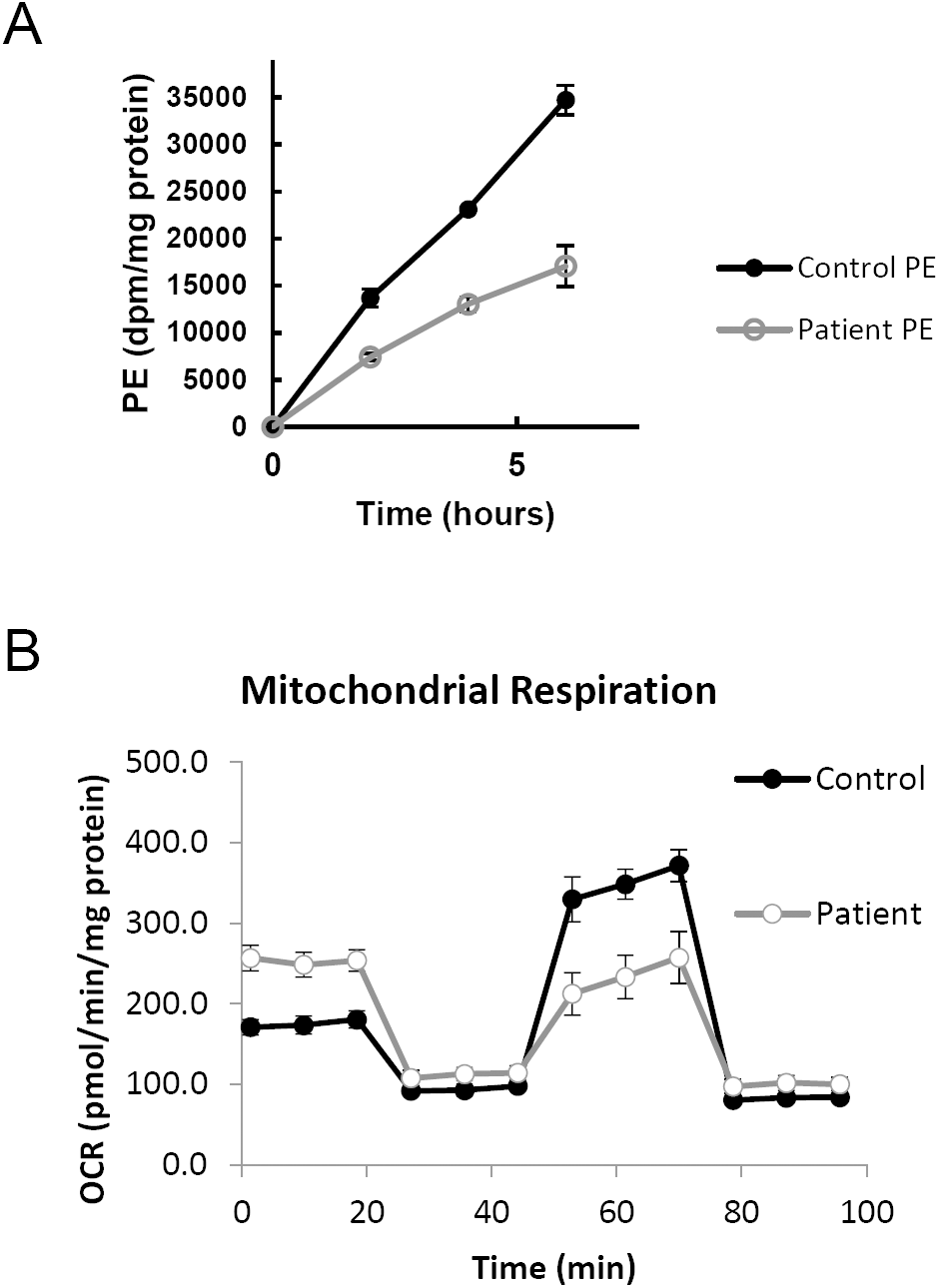
Characterization of PE synthesis and mitochondrial function in patient fibroblast cells. **A)** Patient fibroblasts have decreased ability to convert PS into PE. Control and patient fibroblasts were labeled with [^3^H]serine for 2, 4 and 6 hours, after which total PE was isolated by thin layer chromatography, and the incorporation rate of the [^3^H] label into PE was quantified. **B)** Patient fibroblasts have decreased maximal respiration. Profiles of oxygen consumption rate (OCR) for control and patient fibroblast cells, measured using the Seahorse Extracellular Flux XF24 Analyzer.

Next, we sought to investigate the effect that PISD deficiency might have on mitochondrial function. We observed marked differences between control and PISD patient fibroblasts using a Seahorse metabolic flux analyzer. While basal respiration was elevated in PISD fibroblast cells, maximal respiration was decreased. In fact, PISD fibroblasts had no reserve capacity (Fig 2B). These findings clearly demonstrate alterations in the mitochondrial activity of PISD fibroblast cells.

To look further at mitochondrial function, we quantified mitochondrial morphology via confocal microscopy, as mitochondrial fragmentation is associated with mitochondrial dysfunction. In particular, PE is considered to be a pro-fusion lipid, and mitochondrial fragmentation has been reported in cells deficient in PISD activity (Steenbergen et al., 2005, Tasseva et al., 2013). A slight shift towards more fragmented mitochondrial morphology was observed in patient fibroblasts cells under normal growth conditions (Fig 3A & B). We then treated cells with 2-deoxyglucose (2DG), which blocks glycolysis and forces cells to rely on mitochondrial function for their energy demands. Notably, this treatment has been used previously to exacerbate fragmentation in multiple mitochondrial disease patient fibroblasts (Guillery, Malka et al., 2008). Upon 2DG treatment, patient fibroblasts exhibited a much more dramatic fragmentation phenotype compared to control fibroblasts (Fig 3B).

**Figure 3.**
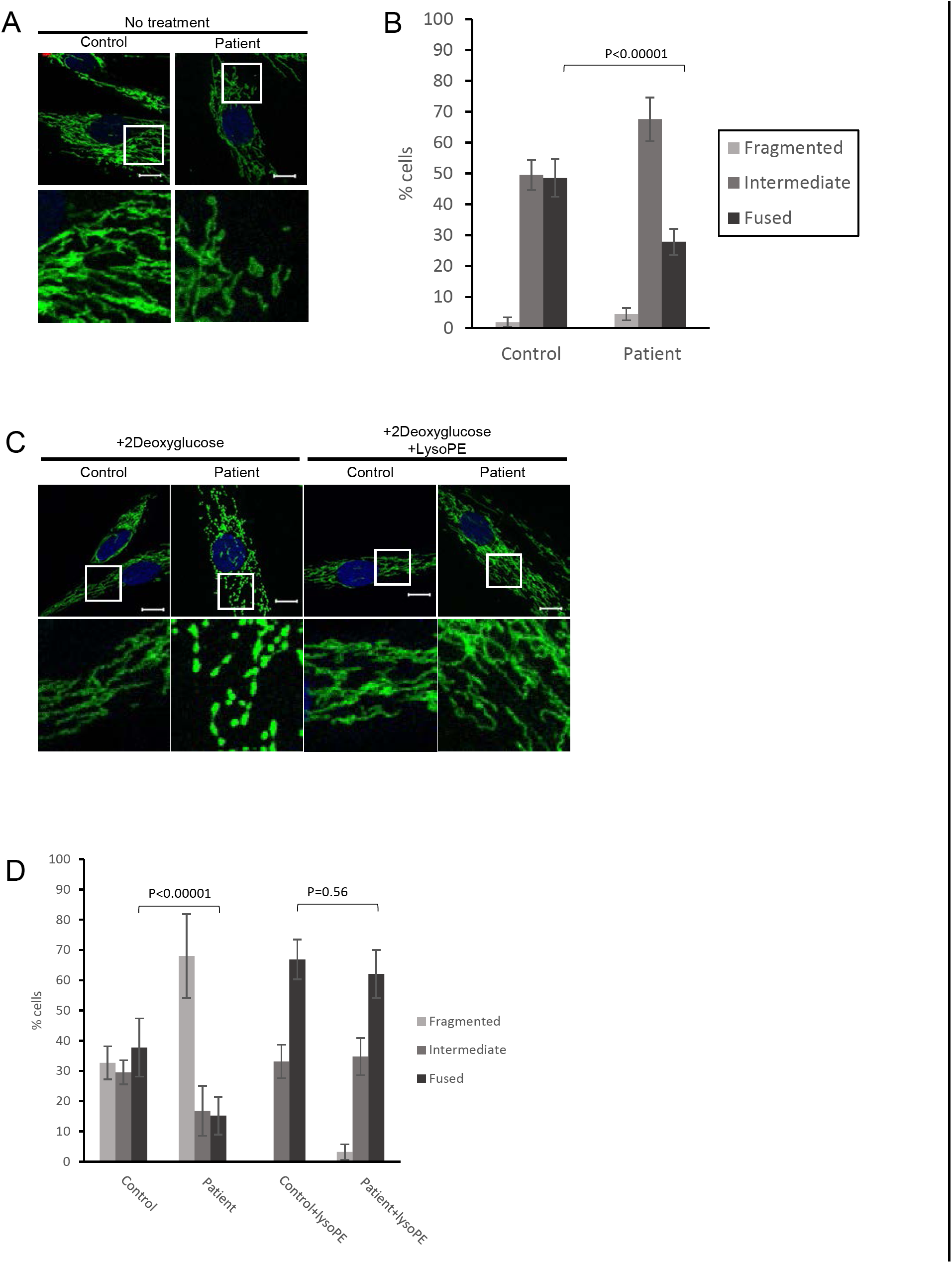
Mitochondrial fragmentation in PISD patient fibroblasts is rescued by supplementation with lyso-PE. **A**) Representative images of mitochondrial morphology under normal growth conditions, stained by immunofluorescence using a TOMM20 antibody (green). Nuclei were stained with DAPI (blue). Bottom panels are a magnification of the white boxed areas shown in the upper panel. Scale bars indicate 10 μm. **B)** Mitochondrial morphology of cells from A) was quantified in at least 50 cells from 3 biological replicates. P-values were determined by Student‘s T-test compared to the number of fused cells in control. **C)** Representative images of mitochondrial morphology of cells treated with 2-deoxyglucose (20 mM) or lyso-PE (50 μM) for 48 hours, as indicated. **D)** Quantification of mitochondrial morphology of cells from (C).

We also examined lysosome structure, which can also be impaired by mitochondrial dysfunction (Demers-Lamarche, Guillebaud et al., 2016). We observed a marked increase in the number of PISD patient fibroblast cells exhibiting enlarged lysosomal structures compared to control fibroblast cells (Fig 4). Taken together with the alterations in mitochondrial function and morphology, these findings provide strong evidence of mitochondrial dysfunction in PISD patient fibroblasts.

**Figure 4.**
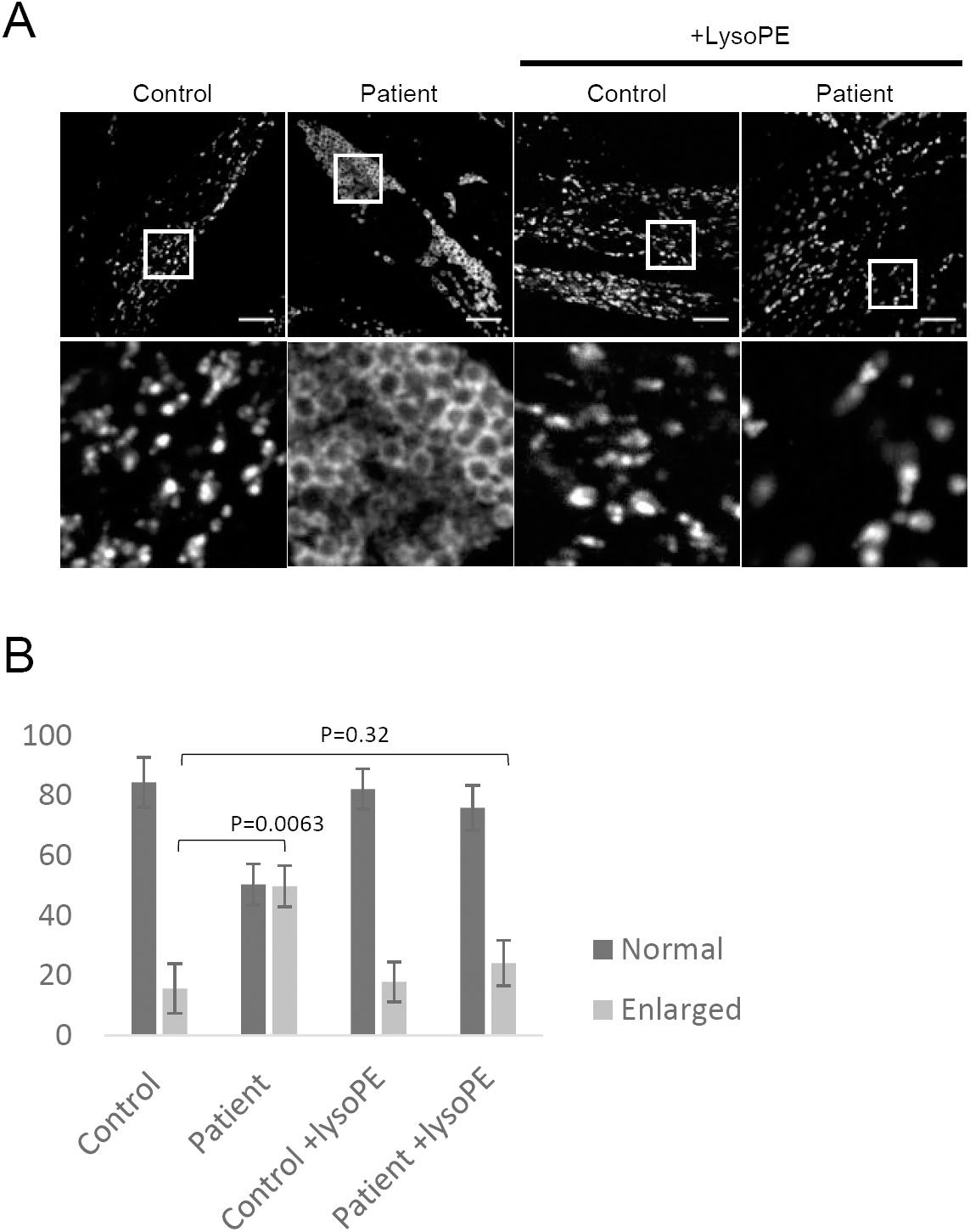
Altered lysosomal morphology in PISD patient fibroblasts is rescued by supplementation with lyso-PE. **A)** Representative images of lysosomes stained by immunofluorescence (IF) against LAMP1 (orange). Cells were grown in either normal medium or medium supplemented with lyso-PE (50 μM) for 48 hours, as indicated. Bottom panels are a magnifications of the white boxed areas in the upper panel. Scale bars indicate 10 μm. **B)** Quantification of lysosomal morphology from three independent experiments, each counting at least 100 cells per condition. P-values were determined by Student’s T-test compared to the number cells with enlarged lysosomes in control.

Finally, in order to link the mitochondrial dysfunction in PISD patient fibroblasts to deficient PISD activity, we treated cells with lyso-PE, which can replenish the mitochondrial pool of PE when PISD activity is impaired (Tasseva et al., 2013). While lyso-PE treatment promoted mitochondrial fusion to some degree even in control cells, we note that the extreme fragmentation observed in patient cells treated with 2DG was completely rescued via lyso-PE supplementation (Fig 3B). Moreover, lyso-PE treatment also restored lysosomal morphology (Fig 4). These findings provide evidence that impaired mitochondrial morphology in the patient fibroblasts is due to low levels of PE in the cells, and that the mitochondrial dysfunction was causing secondary effects on lysosome structure.

### Functional Characterization of PISD Mutations

Additional functional analyses were performed to provide proof that the two PISD variants in our patients were pathogenic. First, we examined the effect of the c.697+5G>A variant on splicing of the *PISD* mRNA. This variant located in intron 5, five base pairs after the end of exon 5, is predicted to weaken the endogenous splice donor site at the end of exon five, with a cryptic splice donor site located 76 base pairs upstream predicted to occur instead. The use of the upstream cryptic splice site would result in the skipping of the end of exon five, with a 1 base pair frameshift for 51 amino acids, and ending in a premature stop codon in exon seven. To investigate *PISD* splicing products, we generated cDNA from control and patient fibroblast RNA. PCR amplification across the region containing the splice sites allowed us to visualize an alternative splicing product in patient samples as a minor PCR band that was not present in control (Fig 5). We also note that treating cells with cycloheximide increased the abundance of the mis-spliced transcript, demonstrating that it is subject to nonsense-mediated decay.Additionally, when performing quantitative RT-PCR across cryptic splice donor site, distinct peaks were observed in the melt-curves from control and patient samples (Sup Fig 1A). Finally, a similar pattern of mis-splicing was observed in cDNA generated from RNA peripheral blood samples provided by patient II-2 and the mother (I-2), as a low level signal in the sequencing electropherograms (Sup Fig 1B). Together, these findings indicate that the c.697+5G>A mutation does indeed alter splicing.

**Figure 5.**
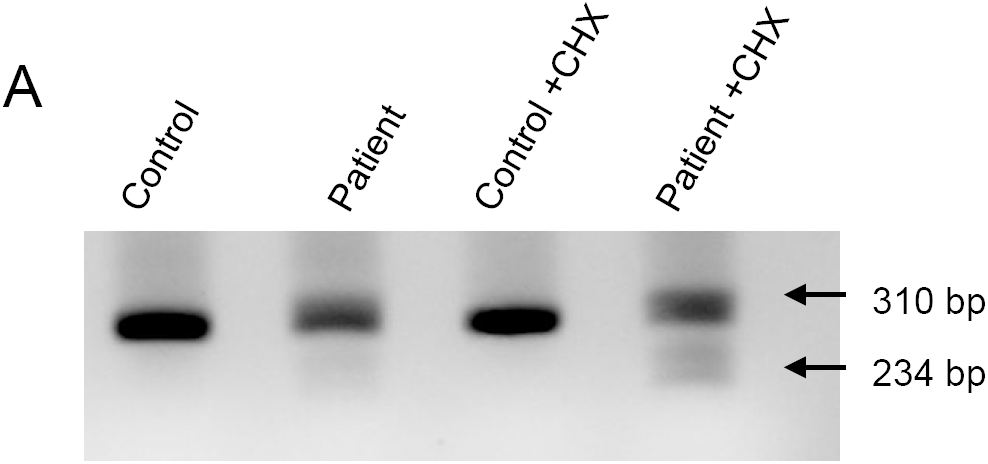
Evidence for alternative splicing and nonsense mediated decay (NMD) of PISD mRNA in patient fibroblasts. **A)** cDNA generated from control and patient fibroblasts was amplified by PCR across the region containing the splice sites in question. Cells were treated with cycloheximide (CHX) (10 μg/ml for 4 hours) to block nonsense-mediated decay, as indicated. The 310 bp product corresponds to the normal splice product. The smaller ~234 bp product corresponds to the predicted alternative splicing product, is visible in patient cDNA, and is more abundant following CHX treatment.

Next, we characterized the c.830G>A [p.Arg277Gln] mutation, which was predicted to be pathogenic. To begin, we utilized the yeast *Saccharomyces cerevisiae*, an established and robust model for mitochondrial disease (Baile & Claypool, 2013). Previously, a protein sequence alignment of phosphatidylserine decarboxylase (PSD) enzymes from humans to bacteria was used to identify evolutionarily conserved residues that are essential for the autocatalytic proteolysis of yeast phosphatidylserine decarboxylase 1 (Psd1p) (Ogunbona, Onguka et al., 2017). Interestingly, R358 in yeast Psd1p, corresponding to human R277, occurs two residues downstream of one of the four missense mutations present in a *temperature sensitive PSD1* allele (Figure 1C) (Birner, Nebauer et al., 2003). Further, it is also in the vicinity of His345, a component of the conserved catalytic triad that is essential for Psd1p autocatalytic proteolysis (Ogunbona et al., 2017). As such, we hypothesized that the R358Q mutation may disrupt Psd1p autocatalytic proteolysis via a mechanism that is enhanced at elevated temperatures. Consistent with this hypothesis, autocatalytic proteolysis of the R358Q mutant, which was relatively normal at 30°C, was significantly impaired at 37°C (Figure 6B). In contrast, self-processing of WT Psd1p into separate α and β subunits occurred at both tested temperatures. While autocatalytic proteolysis was impaired for the R358Q mutant at elevated temperature, it was not completely ablated. Consistent with the persistence of some functional enzyme, the R358Q mutant enabled growth of the *psd1*Δ*psd2*Δ strain at 30°C and 37°C in the absence of ethanolamine (Figure 6C). These results establish the R358Q mutant as a novel *temperature-sensitive PSD1* allele with severely impaired, but not completely ablated, autocatalytic proteolysis at super-physiological temperatures.

**Figure 6.**
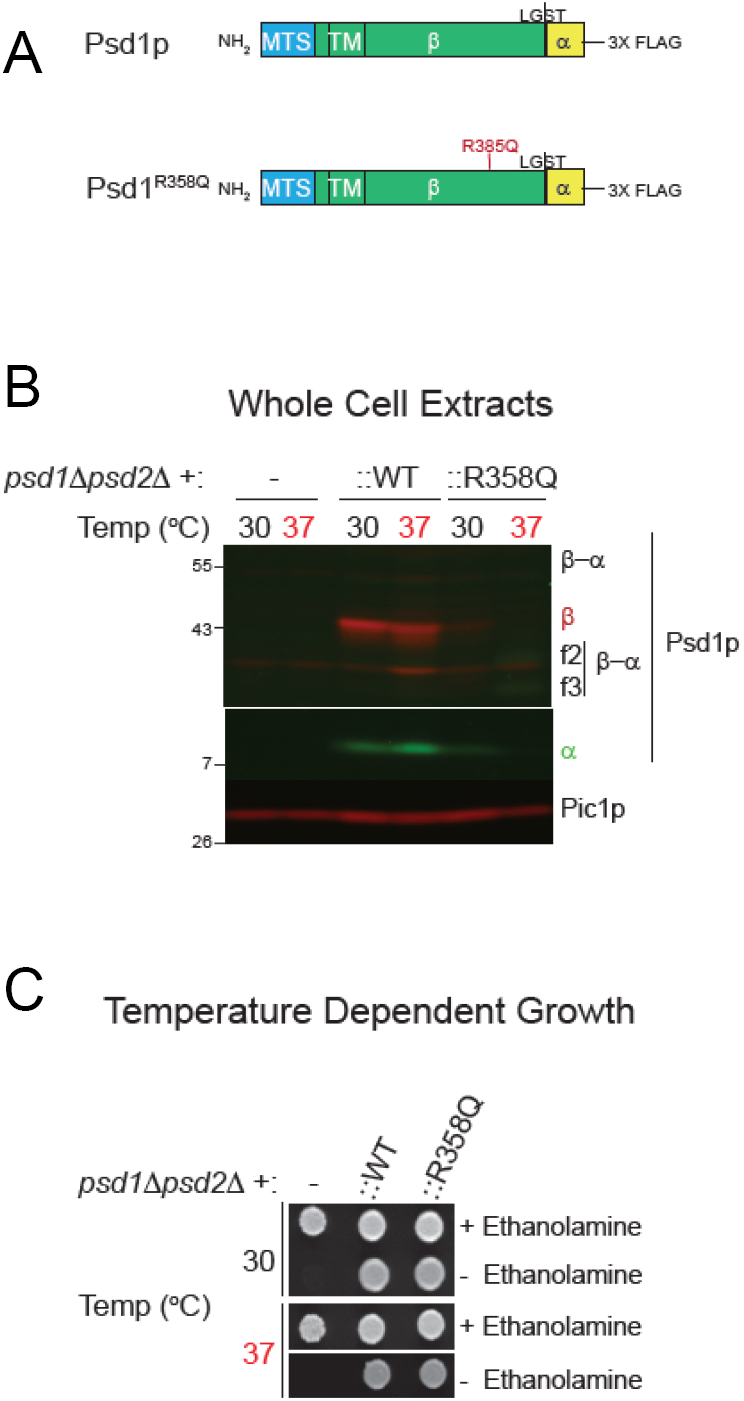
The R277Q mutation disturbs autocatalytic proteolysis when modeled in yeast Psd1p. **A)** Schematic of Psd1p constructs. The patient mutation is indicated. MTS, mitochondrial targeting signal; TM, transmembrane domain. A 3XFLAG tag was added to the C-terminus of both constructs to enable detection of the α subunit post-autocatalysis with anti-FLAG antibodies. **B)** The α (anti-FLAG mouse monoclonal) and β (anti-Psd1p rabbit antisera) subunits of Psd1p were analyzed by immunoblotting in cell extracts isolated from cultures grown at the indicated temperatures; Pic1p served as a loading control. α-β, indicates Psd1p that has not undergone autocatalytic proteolysis. f2 and f3 mark proteolytic fragments generated from the non-processed Psd1p precursor. The migration of molecular mass markers in kDa is indicated at the left. **C)** Pre-cultures (30°C) of *psd1*Δ*psd2*Δ yeast, untransformed or transformed as indicated, were spotted onto SCD plates with or without 2 mM ethanolamine and incubated at 30°C or 37 °Cfor 3 days.

These findings in yeast prompted us to investigate the autocatalytic proteolysis of the human PISD protein containing the R277Q mutation. However, our PISD antibody was unable to detect the endogenous PISD protein in either control or patient fibroblast cells. Thus, we turned to an overexpression approach, where we expressed WT or mutant PISD containing a C-terminal FLAG tag in HEK cells. Following a 48 hour transient transfection of the wild-type PISD, we detected both the 12 kDa alpha and 30 kDa beta PISD subunits with our PISD antibody and a FLAG antibody, respectively. However, we could not detect any of the R277Q mutant protein 24 hour after transfection (Supplemental Figure S2), despite similar levels of *PISD* mRNA expression from the plasmid constructs (Fig 7C). Thus, we looked at protein levels 96 hours following transfection. In addition to the alpha and beta subunits, we also detected a significant accumulation of the unprocessed 45 kDa precursor for the wild-type PISD at 48 hours. In contrast, when the R277Q mutant was overexpressed, only the 45 kDa PISD precursor was observed, indicating that the protein was indeed translated. Thus, similar to experiments in yeast, we note that the R277Q mutation severely impaired the autocatalytic processing of the human PISD protein. As a control, we also generated a PISD construct containing the S378A mutation, which corresponds to the yeast Psd1 S463A mutation that also has impaired autocatalytic processing. Similar to the patient R277Q mutation, when the S378A mutant was overexpressed, the unprocessed 45 kDa PISD precursor was the predominant form.

**Figure 7.**
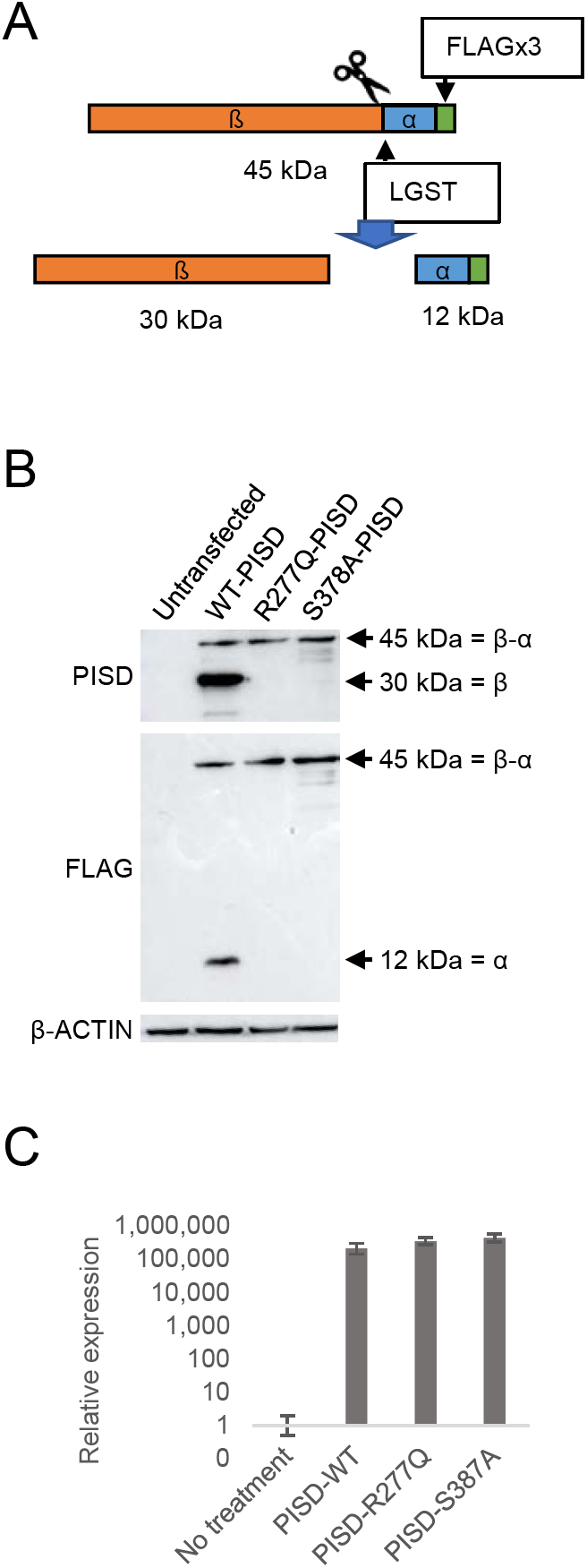
Impaired autocatalytic proteolysis of the R277Q mutation of PISD. **A)** Diagram showing the FLAG-tagged PISD proenzyme that is processed into α and β subunits through autocatalytic cleavage at the conserved LGST site. **B)** Western blot analysis of PISD fragments in HEK cells overexpressing the indicated PISD constructs. Wild type PISD undergoes autocatalytic cleavage into 30 kDa beta-subunits and 12 kDa alpha-subunits required to form an active PISD enzyme. The unprocessed 45 kDa PISD protein is the predominant protein form for the PISD mutant constructs. **C)** Relative expression of PISD mRNAs in HEK cells overexpressing the indicated PISD constructs demonstrates that mutant transcripts are generated.

### PISD and alterations to mitochondrial protein homeostasis

Finally, given that the phenotypic characteristics of our PISD patients resembled those of mitochondrial chaperonopathies, we investigated whether or not mitochondrial proteases as well as mitochondrial protein homeostasis were impaired in PISD patient fibroblast cells treated with 2DG, where we saw dramatic changes in mitochondrial morphology. Given the importance of PE in the IMM we started with OMA1, a zinc metalloprotease that is activated upon mitochondrial stress (Levytskyy, Bohovych et al., 2017). We found that the levels of OMA1 protein expression were severely decreased in PISD patient fibroblast cells (Fig 8A). In order to determine whether this depletion was sufficient to alter protein homeostasis, we examined the cleavage PGAM5, which is processed from a long to a short form by several membrane-bound IMM proteases, including OMA1 (Wai, Saita et al., 2016). In particular, we found decreased levels of the short form of the PGAM5 protein in patient fibroblast cells, suggestive of decreased protease activity. In order to look for more global effects on mitochondrial IMM proteases, we also examined other established targets of mitochondrial IMM proteases. We saw changes to the levels of OPA1, an IMM protein that regulates mitochondrial fusion, and which is regulated by multiple proteases (MacVicar & Langer, 2016), including OMA1 (Ehses, Raschke et al., 2009, Head, Griparic et al., 2009). In addition, we saw decreased levels of the mature form of MRPL32, a mitochondrial ribosomal protein subunit that is processed by the IMM *m-*AAA protease (Nolden, Ehses et al., 2005). Finally, we found that treating PISD patient fibroblasts with lyso-PE rescued the changes in OMA1, PGAM5, OPA1, and MRPL32 proteins.

**Figure 8.**
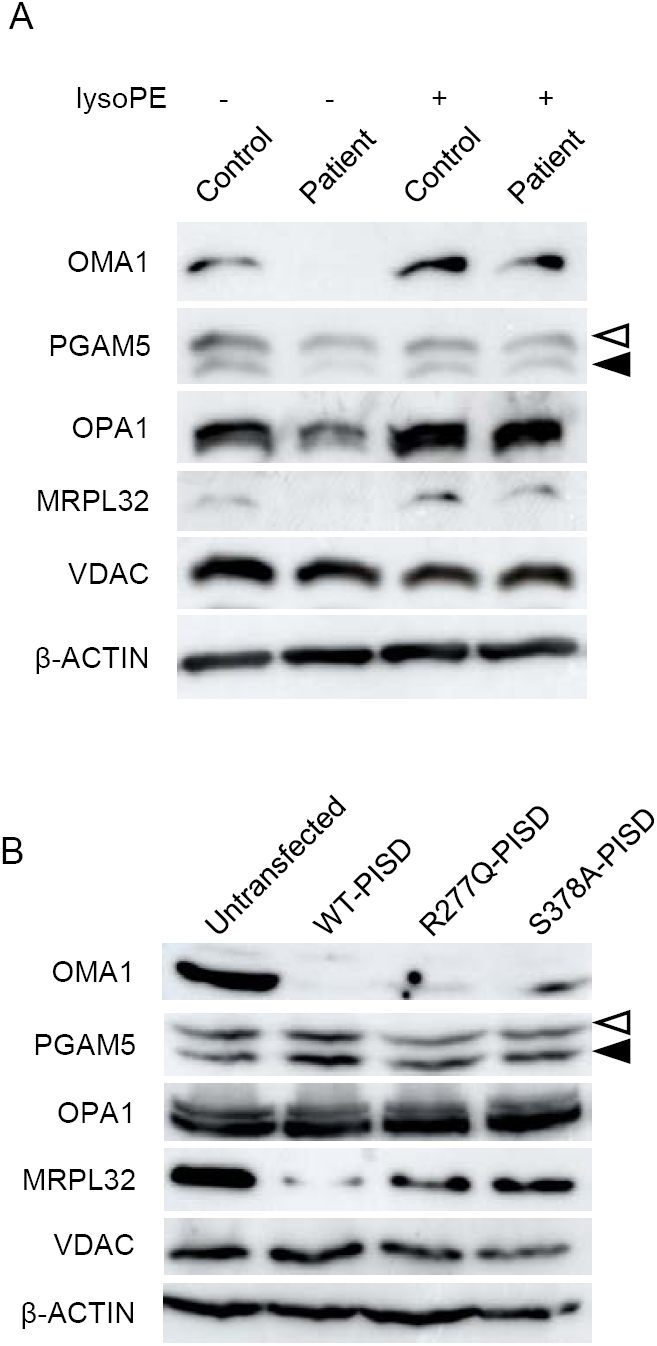
Altered mitochondrial protein homeostasis in PISD patient fibroblasts. **A)** Western blot analysis of various mitochondrial proteins in control and patient fibroblasts treated with 2-deoxyglucose (20 mM) and 50nM lyso-PE for 48 hours as indicated. VDAC was used as a marker for total mitochondrial signal, while β-ACTIN was used as a general load control. Open and solid triangles indicate unprocessed and processed forms of PGAM5. Decreased levels of OMA1, OPA1, MRPL32, and processed PGAM5 were observed in patient fibroblast cells, but rescued upon lyso-PE treatment. **B)** Protein extracts from HEK cells overexpressing wild-type or mutant PISD constructs (as in Fig 7B) were analyzed by Western blot for the same proteins as in part A). Overexpression of wild-type PISD leads to a dramatic decreased in OMA1 and MRPL32, which is blunted when mutant PISD proteins are overexpressed.

We also examined the effects of overexpressing wild-type and mutant PISD on mitochondrial protein homeostasis in HEK cells (Fig 8B). Notably, overexpressing WT-PISD alone significantly reduced the levels of OMA1 and MRPL32, but did not have any major effects on PGAM5 or OPA1. Meanwhile, overexpressing the R277Q and S378A PISD mutants also decreased levels of OMA1 and MRPL32 proteins, although to a less extent than WT-PISD. Altogether, these data are consistent with the notion that altering levels of PE in the IMM, either up or down, impacts mitochondrial proteases and alters mitochondrial protein homeostasis.

## Discussion

Consistent with impaired PISD activity, our characterization of PISD patient fibroblasts revealed impaired phospholipid metabolism and mitochondrial dysfunction, which was exacerbated by 2DG. The fact that supplementation of patient fibroblasts with lyso-PE rescued mitochondrial fragmentation and impaired mitochondrial protein homeostasis in 2DG-treated patient cells strongly supports a mitochondrial PE deficiency in these cells. Furthermore, functional characterization of the PISD patient mutations shows the presence of an alternative splicing event of the maternally encoded transcript, and reduced autocatalytic proteolysis of the paternally encoded protein, demonstrating that these polymorphisms reduce PISD activity and are pathogenic.

An initial quandary when PISD was identified as the only candidate gene, was why the patient phenotype doesn’t resemble classic mitochondrial disease. However, the similarity of skeletal phenotypes observed in PISD patients with recently described skeletal abnormalities in mitochondrial chaperonopathies were notable. To this end, our data showing that IMM proteases and mitochondrial protein homeostasis are impaired in PISD patient fibroblasts explains the similar patient phenotypes.

We considered two possible mechanisms by which mutations in PISD might impair mitochondrial protein homeostasis. The first possibility was that accumulation of the unprocessed PISD protein reduced the ability of mitochondrial proteases to maintain normal mitochondrial proteostasis. To this point, it is likely that unprocessed human PISD is degraded by mitochondrial proteases, similar to the situation in yeast, where the unprocessed autocatalytic mutant Psd1 protein is degraded by the OMA1 protease (Ogunbona et al., 2017). In this regard, the fact that the unprocessed PISD protein was only visible 96 hours post transfection is consistent with an inability of mitochondrial proteases to efficiently remove the accumulated 45 kDa precursor. However, given that transcript levels were increased by several orders of magnitude in HEK cells, we are hesitant to draw conclusions on whether normal levels of the mutant PISD protein would be sufficient to impair mitochondrial protein homeostasis, and lead to pathology. Moreover, given that the unaffected father presumably carries a single copy of R277Q, it is unlikely that the presence of the mutant protein alone, at physiological levels, is sufficient to induce pathology. However, as DNA from the father was unavailable, we could not completely exclude the unlike possibility of gonadal mosaicism.

The second, more likely possibility is that the global reduction of PISD activity and subsequent decrease in PE levels, contributes indirectly to impaired mitochondrial protein homeostasis. The fact that depletion of PISD leads to mitochondrial fragmentation and decreased oxidative phosphorylation (Steenbergen et al., 2005, Tasseva et al., 2013) demonstrates that depletion of mitochondrial PE is sufficient to cause cellular defects. To this end, several mitochondrial proteases are associated with, or are integral to, the IMM (*e.g.* HTRA2/OMI, iAAA, mAAA, OMA1, PARL) and their functions can be regulated by membrane lipid domains (Quiros et al., 2015). As such, these proteases are likely sensitive to alterations in membrane lipid composition such as decreased levels of PE.

The relationship among mitochondrial proteases is complex, as they often cleave and activate each other (Levytskyy et al., 2017), and it is not possible to directly measure protease activity. Nonetheless, our Western analysis of mitochondrial proteins in PISD patient fibroblasts provide strong evidence for decreased mitochondrial protease activity and subsequent impairments in mitochondrial protein homeostasis. In particular, we observed a marked decrease in the protein levels of the protease OMA1. This decrease in OMA1 might be due to destabilization of the protein itself because of altered lipid composition and/or increased degradation of OMA1 by other mitochondrial proteases that cleave OMA1. Notably, *oma1*^*-/-*^ mice are viable, and though they have an obesity phenotype and defective thermogenesis (Quiros, Ramsay et al., 2012), it does not match the phenotypes of our PISD patients. This dissimilarity in phenotypes suggests that loss of OMA1 alone is not sufficient to induce the pathology seen in our PISD patients, where there must be additional contributing factors.Certainly, our data point to the fact that multiple IMM proteases are affected in PISD fibroblasts, as we also see differences in OPA1 and MRPL32 protein profiles. Importantly, the rescue of these protein profiles in patient fibroblasts supplemented with lyso-PE demonstrates that the altered mitochondrial protein homeostasis is due to depleted PE levels. Finally, the fact that we also see changes to mitochondrial protein homeostasis when PISD is overexpressed, albeit with a different profile than the changes in PISD fibroblasts, demonstrates that higher levels of PE can also be detrimental. The different mitochondrial protein profiles in patient fibroblasts deficient in PE and HEK cells overexpressing PISD likely reflects that the various IMM proteases respond differently to either an excess or deficiency of PE levels. Altogether, our data support the notion that IMM proteases are sensitive to changes in PE levels.

Another open question pertains to the loss of PISD activity required for pathogenesis The fact that both parents were unaffected, combined with the fact PISD +/-mice appear normal (Steenbergen et al., 2005), suggests that the threshold of PISD depletion required for pathogenicity lies somewhere between 50% and 100%. In this light, it is likely that the PISD patients still produce some functional PISD protein, as PISD knockout mice are embryonic lethal in mice (Steenbergen et al., 2005). Although we cannot currently assign the relative contribution of each patient allele to the remaining PISD function in patients, it is notable that the R358Q mutation in yeast Psd1 retains some activity. We also note that overexpression of the R277Q and S378A mutants in HEK cells, led to a milder depletion of OMA1 and MRPL32 compared to control, consistent with reduced activity. Moreover, it is also possible that a portion of the maternal allele is spliced normally, or that the amount of alternative splicing varies in different tissue types.

Collectively, our data demonstrate that mutations in *PISD* lead to mitochondrial dysfunction that likely leads to the unique array of patient phenotypes, and demonstrate that *PISD* is a novel human disease gene. Although cataracts are often observed in classic mitochondrial disease, the severe growth impairment with non-specific skeletal anomalies and dysmorphic features are not. However, the combination of skeletal abnormalities with cataracts, distinctive craniofacial features including depressed nasal ridge and intellectual disability are all reminiscent of patient phenotypes described in the recently described subclass of mitochondrial chaperonopathies. Our data provide evidence for impaired mitochondrial protein homeostasis as a result of *PISD* mutations, which likely explains the similarities of the PISD patient phenotype to mitochondrial chaperonopathies. Thus, the addition of *PISD* to the list of genes mutated in mitochondrial chaperonopathies will help define a better mechanistic understanding of this recently appreciated class of mitochondrial disease. Furthermore, this work adds to the broad phenotypic spectrum in mitochondrial phospholipid metabolism-based diseases (Lu & Claypool, 2015).

## Methods

### Patient selection and clinical investigations

The two patients were ascertained through the clinical practice of one of the authors at the Alberta Children’s Hospital. Consent for participation from both patients and their mother was obtained as part of a study approved by the Conjoint Ethics Review Board of the University of Calgary. Investigations such as basic genetic and metabolic blood work, skeletal surveys and MRI scans on both patients had been obtained during the course of their routine clinical care. As they were undiagnosed after conventional investigations they were enrolled in a whole exome sequencing based research study to identify the cause of their presumed genetic rare disease.

### Molecular genetics studies

Genomic DNA was extracted from peripheral blood provided by the patients and their unaffected mother using an Autopure LS system and Gentra Puregene blood kit (Qiagen, Toronto, ON). The Agilent SureSelect V5 All ExonKit (Agilent, Santa Clara, California, USA) and Illumina HiSeq 2500 (San Diego, California) were used for target enrichment and sequencing, respectively. Read alignment to the hg19 reference genome, as well as variant calling and annotation, were completed as described in previously published Care4Rare Consortium projects (Beaulieu, Majewski et al., 2014).

Due to the rarity of the sisters’ phenotype, annotated variants with an ExAC minor allele frequency less than 1% and seen in less than five previous Care4Rare samples were considered. An autosomal recessive mode of inheritance was assumed and included analysis of: rare homozygous variants shared by both sisters and carried by the mother; or, rare bi-allelic variants shared by the sisters, of which only one of the variants was carried by the mother. Resulting rare variants and their respective genes were visualized using Alamut^®^ Visual version 2.9.0 (Interactive Biosoftware). Conservation, *in silico* pathogenicity prediction scores (Polyphen, SIFT and MutationTaster), and splice prediction scores (SpliceSiteFinder-like, MaxEntScan, NNSPLICE, GeneSplicer and Human Splicing Finder) were taken in account. Sanger sequencing, using the ABI BigDye Terminator Cycle SequencingKit v1.1 (Life Technologies, Carlsbad, California, USA) on a 3130xl genetic analyzer (Life Technologies), confirmed the candidate variants.

### cDNA Sequencing

A PAXgene Blood RNA kit (PreAnalytiX) was used to extract RNA from peripheral blood samples provided by patient II-2 and their mother. Complementary DNA (cDNA) was synthesized (SuperScript III, Invitrogen, Carlsbad, California, USA) from the extracted RNA. PCR amplification (HotStar Taq Plus, Qjagen) across exon junctions of cDNA was performed using primers 5’-GGCGAAATGGTTGCACTTC-3’ (forward) and 5’-GCAGGTCCCGGTCAAAG-3’ (reverse). Patient cDNA was Sanger sequenced using the 5’-GGCGAAATGGTTGCACTTC-3’ (forward) and 5‘;-GCAGGTCCCGGTCAAAG-3’ (reverse) primers as well as a sequencing primer 5’-GATGGAGGCCCGTAAGC-3’ (forward) to ensure complete capture of exons five through eight of *PISD* (Ref Seq NM_014338.3).

### Fibroblast cell culture growth conditions and treatments

Fibroblast cells were obtained from patient-and gender/age-matched control from skin punch biopsies. Fibroblast cells were cultured in minimum essential medium (MEM), without glutamine (Gibco) but supplemented with 2 mM of L-glutamine (Gibco) and 10% heat inactivated FBS (Gibco). Fibroblast cells were detached from 10 cm culture dishes (Fisher) using Trypsin-EDTA (0.25%) and passed in a 1:3 ratio 3 times per week. For confocal microscopy experiments, Fibroblast cells were seeded at 20,000 cells per well using 24 well plates on glass cover slips (12-545-81 Fisher Scientific) overnight. Fibroblast cells were treated with 50 nM lyso-PE (Sigma-Aldrich 89576-29-4) in ethanol and/or 20 mM 2-Deoxyglucose (Santa Cruz sc-202010) for 48 hours.

### HEK cell culture and transfection

HEK cells were cultured in Dulbecco’s modified Eagle’s medium (DMEM) (Gibco) with 10% heat-inactivated FBS (Gibco). Cells were seeded at 2 × 10^6^ cells per plate on 10 cm plates overnight the night before transfection. Cells were transfected using Lipofectamine 3000 (ThermoFischer Scientific) reagent with 15 μg plasmid DNA per plate. The human *PISD* ORF corresponding to Isoform A (NCBI Reference Sequence (NP_001313340.1) was amplified by PCR using cDNA from HeLa cells, cloned initially into pSP64 (Promega), and then subcloned into pcDNA5/FRT (Invitrogen) with a 3XFLAG tag appended to its carboxy-terminus by overlap extension enabling detection of the α subunit. The R277Q and S378A mutant variants were generated by overlap extension PCR.

### Immunofluorescence

Cover slips were fixed in phosphate-buffered saline buffer containing 4% paraformaldehyde at 37°C for 15 min followed by quenching with ammonium chloride (50 μM).Fixed cover slips were stained with an αTOMM20 antibody (sc-11415, Santa Cruz) for detection of mitochondria, or an αLAMP1 antibody (sc-11415, Santa Cruz) for detection of lysosomes. Samples were then incubated with appropriate Alexa Fluor-labelled secondary antibodies (Invitrogen). Finally, cover slips were mounted for imaging with Dako fluorescent mounting medium containing DAPI (S302380, Agilent Technology).

### Mitochondrial morphology

Mitochondrial morphology was imaged using 488 nm lasers with 60x lens on a ZEISS LSM 700 confocal laser-scanning microscope. Quantification of mitochondrial morphology was performed by grading three replicates of at least 50 cells each, into 3 levels of mitochondrial morphology: fused, intermediate and fragmented according to representative images of each level.

### Lysosomal morphology

Lysosomal morphology was imaged using 568 nm lasers with 100x lens on an Olympus SD-OSR imaging system. Quantification of lysosomal morphology was performed by scoring three replicates of 100 cells each, as either normal or enlarged according to representative images of each level.

### Oxygen consumption and respiratory function

Fibroblast were seeded at 40,000 cells per well and grown overnight in XF24 plates (Agilent). On the day of the experiment, normal growth medium was replaced by Seahorse XF Base Medium containing 1 mM pyruvate, 2 mM glutamine, and 10 mM, pH 7.4. After 45 min incubation at 37°C without CO_2_, the cells were loaded into the Seahorse XF24 Analyzer for calibration. Oxygen consumption rate (OCR) was measured at 3 time points following serial incubations with oligomycin (final concentration 0.5 μM), FCCP (final concentration 1.0 μM) and antimycin (final concentration 0.5 μM). At the end of the experiment cells were lysed with radioimmunoprecipitation assay (RIPA) buffer and total protein was determined by bicinchoninic acid (BCA) assay. OCR was normalized to total protein/well.

### Western blotting

Total protein lysates were prepared using RIPA buffer. Protein concentration was determined by BCA assay. Protein (50 μg) from whole cell lysate was boiled with Laemmli sample buffer (Bio-Rad) was boiled and loaded into each lane of 12% SDS-PAGE gels then transferred to PVDF membranes. The membranes were then blocked with 5% skim milk and blotted with various primary antibodies: anti-β-ACTIN (A5316, Sigma), anti-PISD (Custom made), anti-FLAG (Novus Biological, NB600-344), anti-MRPL32 (Assay Biotech, C14071) anti-OMA1 (Abcam, ab104316), anti-OPA1 (BD Bioscience, 612606), anti-PGAM5 (Sigma, HPA036978), and anti-VDAC (Abcam, ab14734). All primary antibodies were used in 1:1000 dilution. Appropriate secondary HRP-conjugated antibodies (ThermoFischer Scientific) were used in 1:10,000 dilution. Custom-made monoclonal antibodies against human PISD were produced by ABclonal Science Inc. using purified recombinant protein as antigen (details below). A total of five hybridomas (PISD-1-3-4-2, PISD-1-6-1-2, PISD-2-3-3-3, PISD-2-6-2-1, and PISD-2-9-2-1) were established that secreted PISD reactive antibodies. Protein G purified monoclonal antibodies from PISD-2-9-2-1 were used in this study.

### Purification of recombinant human PISD

The predicted mature human PISD spanning residues 77-409 was cloned downstream of the 6x His tag provided by the pET28a vector (Novagen). Transformed BL21(DE3) pLysS RIL *E. coli* cells were used to produce recombinant His6PISD. A 1 liter culture of cells was induced at 37 °C with 1 mM IPTG for 4 hours and the cell pellet collected by centrifugation at 3,020 x*g* for 10 min. The cell pellet was resuspended in 250 mL of 0.9% NaCl solution, centrifuged at 3,020 x*g* for 10 min, and the supernatant again discarded. The cell pellet was resuspended in 40 mL of Lysis Buffer (50 mM NaH2PO4, 300 mM NaCl, 10 mM imidazole, 1% Tween-20, 0.1 mM EDTA, pH 8.0) and the cells were incubated with 40 mg/mL lysozyme for 30 min on ice. Cell lysis was accomplished using an Avestin homogenizer. The cell lysate was centrifuged at 10,000 x *g* at 4 °C for 20 minutes and the cell pellet resuspended in Inclusion Body Solubilization Buffer (1.67% Sarkosyl, 0.1 mM EDTA, 10 mM dithiothreitol, 10 mM Tris-HCl, 0.05% PEG, pH 7.4), followed by 20 minutes incubation on ice. The suspension was mixed with 14 mL of 10 mM Tris-HCL (pH 7.4), centrifuged at 12,000 x *g* at 4 °C for 10 min, after which the pellet was discarded. Protein in the supernatant was purified by passing the suspension through a Ni-NTA column (Qiagen) and eluted with Elution Buffer (250 mM imidazole, 0.1% Sarkosyl, 50 mM NaH2PO4, 300 mM NaCl, 10% glycerol, pH 8.0). The purified protein was run on a 12% SDS-PAGE gel, then further isolated from contaminating proteins by removing a gel slice corresponding to 41 kDa and incubating it with 50 mL 1X PBS at 37 °C overnight. The protein was then concentrated by centrifugation with an Amicon Ultra-4 (Millipore) filter at 4,000 x *g* for 10 min. The resulting 4.5 mL of purified protein was dialyzed overnight in 0.5% SDS in PBS at room temperature to remove excess SDS. This resulted in 8.3 mg of His6PISD in 4.5 mL.

### Confirmation of alternative splicing variant

Total cellular RNA was isolated using total RNA kit (Omega) according to the manufacturer’s instructions. cDNA was synthesized by iScript advanced cDNA kit (Bio-Rad) according to manufacturer’s instructions. PCR-amplification of the PISD splice variant was performed using Taq polymerase (Invitrogen) at 58°C for 45 cycles on Bio-Rad S1000 thermal cycler. Template DNA (300 ng) was used with 10 μM of forward and reverse primers. Forward primer 5’-GCCTGCACAGCGTGATTAG −3’ and reverse primer 5’-TGACATCAGGGAGCCTGG −3’ were chosen at exon junctions to eliminate genomic DNA amplification. Resulting PCR product was visualized with EZ vision dye loading buffer (Life Technology) on 1.5% agarose gel electrophoresis

### Yeast Modeling

All yeast strains used were derived from GA74-1A (*MAT***a** *his3-11,15 leu2 ura3 trp1 ade8* [*rho+ mit+*]). The *psd1*Δ*psd2*Δ (*MAT***a** *leu2 ura3 ade8 psd1*Δ::*TRP1 psd2*Δ::*HISMX6*) was described previously (Onguka, Calzada et al., 2015). The *psd1*Δ*psd2*Δ strain expressing WT Psd1p with a 3XFLAG tag on its C-terminus enabled separate detection of the α and β subunits as described (Onguka et al., 2015). The R358Q pathogenic yeast mutant was generated by overlap extension (Ho, Hunt et al., 1989) using pRS305Psd3XFLAG (Onguka et al., 2015) as the template. To generate a *psd1*Δ*psd2*Δ strain expressing the R358Q mutant, *psd1*Δ*psd2*Δ yeast were transformed with the linearized pRS305-based plasmid and genomic integrants selected on synthetic dropout media (0.17% yeast nitrogen base, 0.5% ammonium sulfate,0.2% dropout mixture synthetic–leu, 2% dextrose). To determine if the R385Q mutant was functional, overnight cultures grown in YPD medium were spotted on synthetic complete dextrose (SCD) plates in the absence or presence of 2 mM ethanolamine and grown at the indicated temperature. Preparation of yeast cell extracts and immunoblotting was performed as described (Claypool, McCaffery et al., 2006). Antibodies against the Psd1p β subunit and Pic1p were generated in the Claypool laboratory and described previously (Onguka et al., 2015, Whited, Baile et al., 2013). Other antibodies used were mouse anti-FLAG (clone M2; Sigma) and Dylight conjugated fluorescent secondary antibodies (ThermoFischer Scientific).

## Acknowledgements

The authors would like to thank the study participants and their family. We would also like to acknowledge the contributions of Drs. Ross McLeod, Rebecca Trussell, Graham Boag, Colleen Adams, Carolyn Skov, and Sheila Unger in the clinical care and previous phenotypic characterizations of this family, as well as Ms. Mary Anderson for clinical support. This work was supported by Alberta Children‘s Hospital Foundation (T.S.); the National Institutes of Health (R01GM111548 to S.M.C.); the National Science Foundation Graduate Research Fellowship (DGE1746891 to P.S.). This work was performed under the Care4Rare Canada Consortium funded by Genome Canada, the Canadian Institutes of Health Research, the Ontario Genomics Institute, Ontario Research Fund, Genome Alberta, Genome BC, Genome Quebec, and Children’s Hospital of Eastern Ontario Foundation.

## Figure Legends

**Figure S1. Evidence for alternative splicing A)** Melt curve plot from triplicate quantitative RT-PCR reactions spanning the predicted splice region shows two distinct peaks for RNA samples generated from control (red) or PISD patient (blue) fibroblasts. **B)** Sequencing traces of PISD cDNA generated from blood. Low levels of an alternative sequence corresponding to the alternative splice product were detected in the affected daughter (II-2) and mother (I-2) (boxed in red).

**Figure S2. The R277Q PISD protein is not detected following 24 hours of overexpression.** Western blot analysis for PISD protein fragments in HEK cells overexpressing the indicated PISD constructs. Both the 30kDa α-subunit and 12 kDa β-subunit are visible for the wild type PISD, but no protein is visible for the R277Q mutant.

